# Polypeptide collapse modulation and folding stimulation by GroEL-ES

**DOI:** 10.1101/2020.06.17.157057

**Authors:** Mohsin M. Naqvi, Mario J. Avellaneda, Andrew Roth, Eline J. Koers, Vanda Sunderlikova, Günter Kramer, Hays S. Rye, Sander J. Tans

## Abstract

Unfolded proteins ubiquitously collapse into a compact yet dynamic state^1,2^. While this compaction is pivotal to protein folding^3^, aggregation^4,5^, intrinsic disorder^6^, and phase separation^7^, its role in protein quality control mechanisms remains obscure^8^. Collapse has been characterized mainly for polypeptides that are free in solution, in terms of kinetics, chain expansion, and effect on folding^9,10^. Yet, theory suggests that the solvent-mediated forces driving collapse can be altered near hydrophobic and charged surfaces, which are observed for many proteins including GroEL-ES^11,12^. Notably, while GroEL-ES is the archetypal protein-folding chaperone, its folding mechanism remains unresolved^13,14^. GroEL-ES is proposed to sterically confine polypeptides within its closed chamber^15^, unfold misfolded states^16,17^, or promote folding indirectly by suppressing aggregation^18,19^. Here, using integrated protein manipulation and imaging, we show that GroEL-ES can strengthen the collapse of polypeptide substrates, and hence stimulate folding directly. Strikingly, attractive forces pull substrate chains into the open GroEL cavity -unclosed by GroES-, and hence trigger a gradual compaction and discrete folding transitions, even for slow-folding proteins. This collapse enhancement is strongest in the nucleotide-bound states of GroEL, and is aided by GroES binding to the cavity rim, and by the amphiphilic C-terminal tails at the cavity bottom. Peptides corresponding to these C-termini alone are sufficient to strengthen the collapse. The results show a mechanism that allows folding to be stimulated: by strengthening the collapse, residues are brought together that must contact to fold. The notion that one protein can modulate the collapse of another may be generally important in protein conformation and coacervation control, for systems ranging from the GroEL-ES homologue TRiC/CCT^20^, to the oncogenic c-Myc/Max complex^21^, and the nuclear pore transporter transportin^22^.

## Folding acceleration by GroEL-ES

We first investigated direct folding stimulation, in the absence of aggregation, by following single GroEL-ES substrates in real time using optical tweezers. We tethered individual Maltose Binding Protein (MBP) molecules between polystyrene beads using DNA handles^23,24^, exposed them to relax-wait-stretch cycles by moving the laser beams that trap the beads, and quantified the fraction of cycles *P_c_* showing refolding into core MBP states (Fig. 1a-c). The core MBP fold is the central and major part of MBP that only lacks a number of external alpha-helices, and is rate-limiting in folding^25–27^. With GroEL-ES and ATP also present, we found that *P_c_* increased modestly from 0.7 to 0.85 (Fig. 1d, Extended data Fig. 1), while the unfolding force *F_u_* remained similar (Fig. 1e).

**Figure 1.**
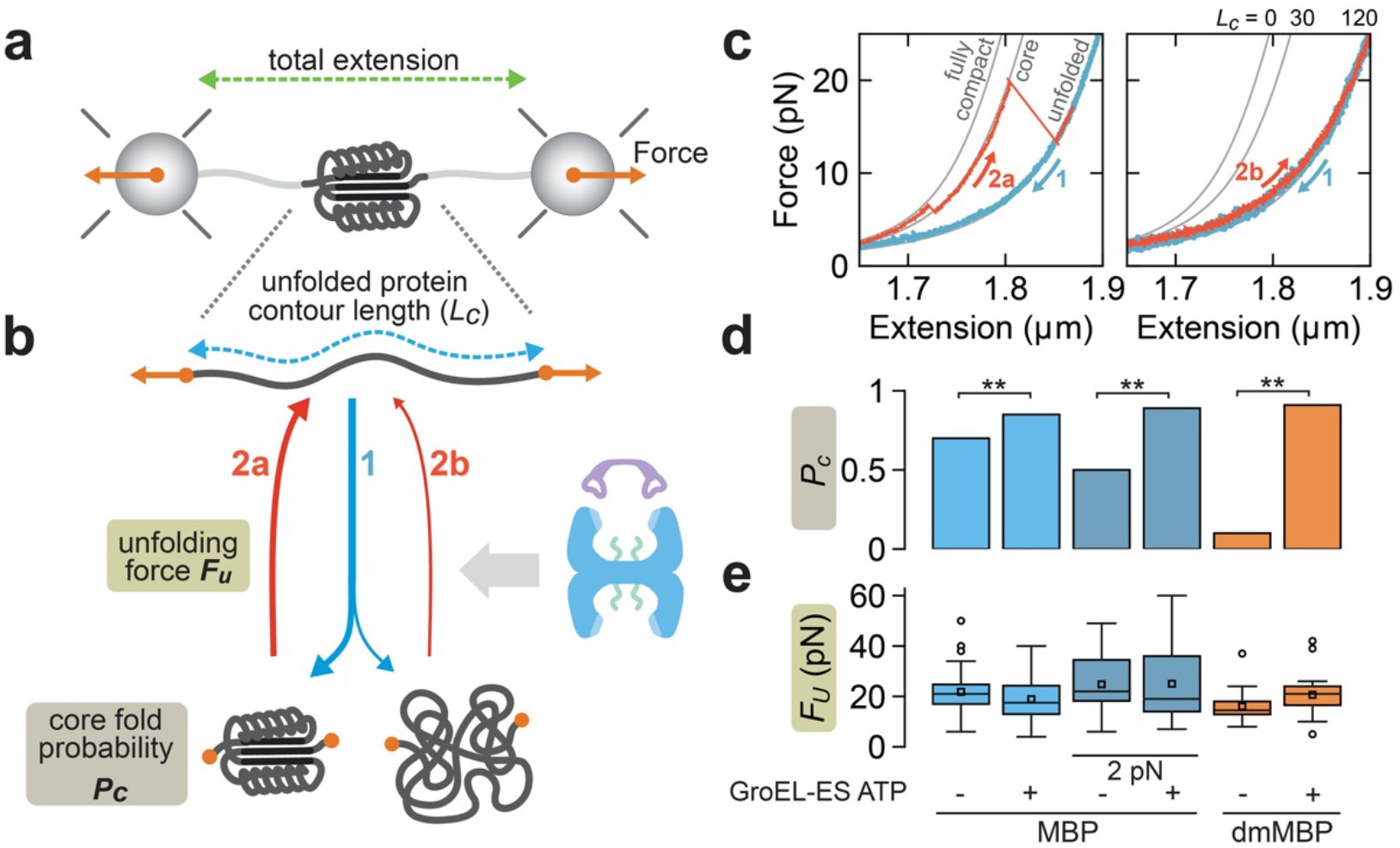
Following single proteins in time shows folding acceleration mediated by GroEL-ES. **a**, Cartoon of optical tweezers experiments. **b**, Relax-wait-stretch cycles to quantify MBP core refolding. Unfolded chains (top) are relaxed, kept at 0 pN for 5 s (bottom), or alternatively at 2 pN for 30 s, and stretched to assess the new state. **c**, Corresponding force-extension data. After relaxation of unfolded MBP (both panels, blue traces, 1), and waiting at 0 pN, stretching data follows the theoretical WLC curve of the MBP core states (left panel, red trace 2a) or the unfolded state (right panel, red trace 2b). Conditions: buffer in absence of GroEL-ES. **d**, Fraction of cycles showing core state refolding (*P_c_*), determined as described in panels b and c. Conditions: alternating with and without 200 nM GroEL, 500 nM GroES, 1 mM ATP, for MBP substrate and waiting at 0 pN (*N*=65 and *N*=53), for MBP at 2 pN (*N*=18 and *N*=21), and for dmMBP at 0 pN (*N*=19 and *N*=21). Two stars: significant difference (p<0.05). (see Extended data Fig. 1). **e**, Force at which the protein fully unfolds (*F_U_*), which is found to be approximately constant for different conditions. When the protein does not unfold, the maximally sustained force is taken. For dmMBP alone, *F_U_* of the first stretching curve is displayed because of the low refolding rate. Conditions as in panel d.

To improve our ability to detect acceleration, the spontaneous folding rate (in the absence of GroEL-ES) was reduced in two ways: 1) Rather than relaxing the chains fully to 0 pN, we maintained a low (about 2 pN) force during the waiting time in the relax-wait-stretch cycles. As a result, the stretched protein chain must now overcome the applied force that counteracts folding, thus limiting the folding rate. 2) We introduced two mutations in MBP (dmMBP) that are known to compromise folding^15^. We found that *P_c_* indeed decreased to 0.5 and 0.06 in these two respective experiments (Fig. 1d). From these lower values in the absence of GroEL-ES, *P_c_* now increased more steeply with GroEL-ES and ATP present (*P_c_* = 0.9, Fig. 1d, Extended data Fig. 1). Overall, these findings showed folding acceleration of single substrates, in the absence of aggregation, while they also indicated that the DNA handles did not inhibit folding stimulation by GroEL-ES. The stimulation mechanism remained unresolved, however.

## Stabilization of substrate chains in their unfolded state

Because of the complex dynamics of the ATP-driven GroEL-ES cycle^13,14,28^, we decided to first examine GroEL in different nucleotide-bound states, in the absence of GroES and ATP hydrolysis. Under these conditions, we now detected a behavior of the MBP chains that was not resolved in the previous experiments. At certain moment during the during relax-wait-stretch cycling, we observed a switch from the regular refolding and unfolding, to a prolonged unfolded state persisting over multiple relax-wait-stretch cycles, until the tether broke (Extended data Fig. 2). This switching occurred most frequently for the APO state (GroEL without nucleotides, 50% of the tethers), and less so for the ATP state (mimicked by the slowly hydrolyzing GroEL398A^29^ and ATP, 30% of the tethers), and the ADP state (GroEL with ADP present, 20% of tethers). These findings are consistent with the known stable binding of unfolded substrates to the apical domains at the rim of the GroEL cavity^30^, and suggested that they can remain bound to extended substrate chains.

## Induced collapse of substrate chains

The above experiments presented other notable features. When tension on a non-interacting polypeptide chain is relaxed, it coils up like a string (Fig. 2a, top). The associated decreases in end-to-end distance (termed extension, Fig. 2b) is then described by the worm-like chain (WLC) model (Fig. 2c-d, left grey curve). However, here the relaxation yielded extensions that became progressively much smaller than expected for a non-interacting polypeptide, decreasing down to values corresponding to a fully compacted polypeptide (Fig. 2d, Extended data Fig. 3a, Extended data Fig. 4b-c, and Extended data Fig. 5a). These data thus indicated a gradual polypeptide compaction process, in which the compacted part of the polypeptide chain increases in size. Conversely, the contour length of the non-compacted part (*L_c_*) thus decreases, as can be quantified using the WLC model (Fig. 2a-b).

**Figure 2.**
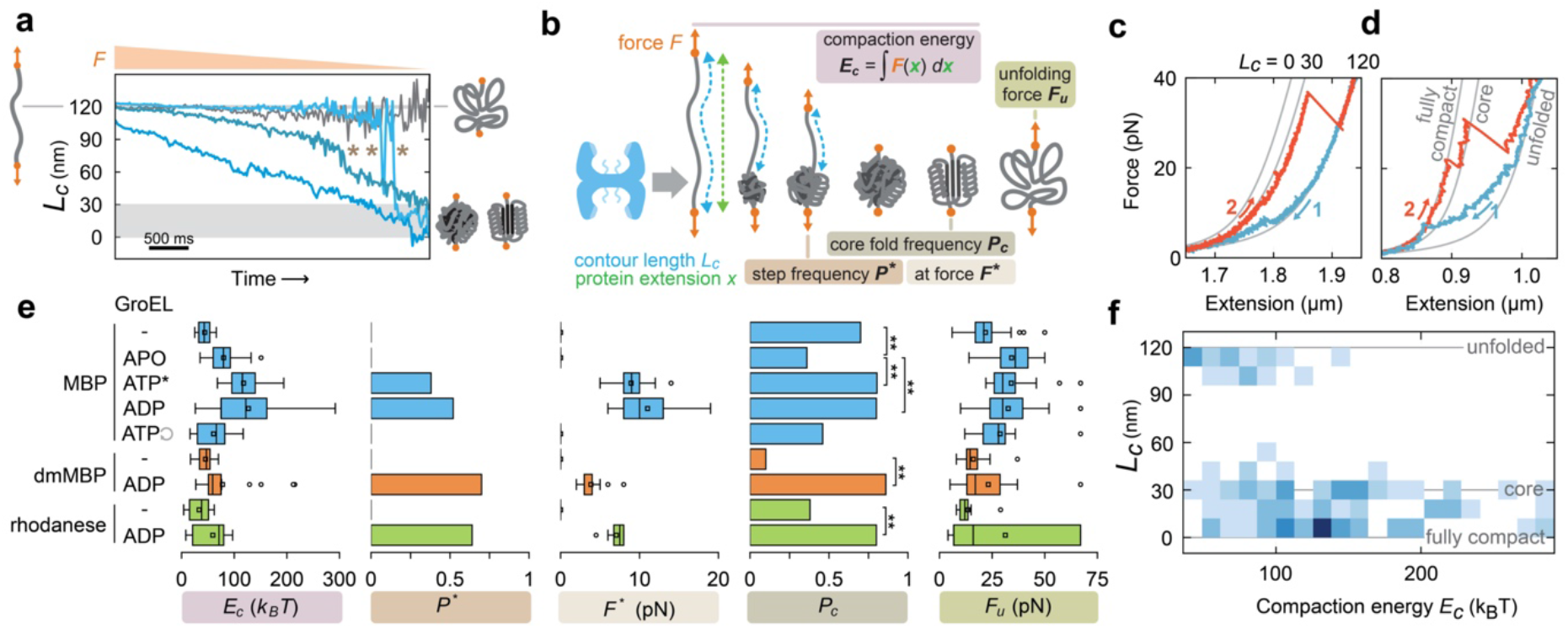
An open GroEL cavity can drive protein chain collapse and folding. **a**, Contour length of the non-compacted part of the chain (*L_c_*) during relaxation, determined using force-extension data (e.g. blue curves panel c and d) and the WLC model. Measurement noise prevents *L_c_* quantification below about 2 pN. Blue curves: dmMBP with 200 nM GroEL and 1mM ADP. Gradual *L_c_* decrease indicates gradual growth of the compacted part of the chain (see panel b). Stars: sudden *L_c_* steps indicating (partial) folding. Gray curve: data illustrating no detectable compaction in accessible force range, with *L_c_* remaining constant as the chain coils (no GroEL). Gray bar: compaction down-to core state or smaller. **b**, Cartoon of relax-wait-stretch cycle and measured quantities. Step frequency *P*\* is the fraction of cycles showing one or more steps during relaxation (panel a, star). The force at which the step occurs is *F*\*. *P_c_* is the fraction of cycles showing refolding to the core state (see Fig. 1 and Extended data Fig.1). The compaction energy *E_c_* is determined using the measured forces and length changes during relaxation, which represent mechanical work (see Extended data Fig. 4). *F_u_* is the force required to fully unfold a protein. **c-d**, Force-extension data of relax-wait-stretch cycles. Relaxation shows gradual compaction despite counteracting applied force, evidenced as a gradual deviation from WLC for non-interacting chain. Conditions: MBP with 200 nM GroEL and 1 mM ADP. Numbers indicate temporal order. **e**, Parameter quantification from relax-wait-stretch cycles in different conditions (see panel b). Conditions: MBP, no chaperones (-, *N*=65), 200 nM GroEL (APO, *N*=33), 200 nM GroEL398 and 1 mM ATP (ATP*, *N*=43), 200 nM GroEL and 1 mM ADP (ADP, *N*=96), 200 nM GroEL and 1 mM ATP (ATP-cyclic arrow, *N*=21), dmMBP no chaperone (-, *N*=19), GroEL-ADP (ADP, *N*=21), rhodanese no chaperones (-, *N*=16), GroEL-ADP (ADP, *N*=20). Two stars: significant difference (*p*<0.05). **f**, *L_c_* of refolded states against *E_c_* of previous relaxation. *L_c_* is defined by the initial stretching data at low force, and the WLC curve it follows. For instance: *L_c_* = 30 (panel c), *L_c_* = 0 (panel d). See also Extended data Fig. 1. Data of MBP with GroEL-ADP and with GroEL398-ATP was combined. The distribution indicates that strong compaction almost always results in a folded state between core and fully compacted. All curves in this latter category showed unfolding via the core state (see panel c and d).

We quantified the compaction strength by extracting the compaction energy *E_c_* for the full relaxation process down to 0 pN, using the measured forces and distance changes that reflect mechanical work (Fig. 2b-c, Extended data Fig. 4). We found that some compaction also occurred for MBP alone, but was stronger with GroEL, in particular in the ATP and ADP states. In principle, such an increased compaction could be the result of the single protein substrate chains binding to multiple GroEL apical domain sites. However, a number of features in the data indicated a different explanation. First, compaction was strongest for the ATP and ADP states, as quantified by both *E_c_* and the fraction *P*\* of traces showing steps (Fig. 2b-c). Yet, the stabilization of unfolded states by apical domain binding is weakest for these nucleotide states, as was discussed in the previous section.

More importantly, the gradual compaction was often accompanied by sudden step-wise compaction events (Fig. 2a, stars). These steps suggested folding transitions rather than stable binding: they were large in size (up to nearly the total chain length, Fig. 2a, Extended data Fig. 3b), occurred at high forces (up to 19 pN, Fig. 2c-e), and exhibited reversible ‘hopping’ transitions characteristic of folding^31–33^ (Fig. 2a, Extended data Fig. 3c). Moreover, after relaxation and waiting, the polypeptides were often observed in the folded MBP core state, as indicated by subsequent stretching data following the core WLC curve (Fig. 2c-f), and seen before for spontaneous and GroEL-ES-assisted folding. Notably, the fraction of cycles showing refolded cores was particularly high for the ATP and ADP states (*P_c_*=0.8 for both, Fig. 2e). *P_c_* increased further beyond 0.95 when the preceding relaxation showed strong compaction (*E_c_*>100 *k_B_*T, Fig. 2f, Extended data Fig. 5). Strong compaction during relaxation of unfolded MBP thus yielded high core folding probabilities. Overall, these findings showed that compaction was distinct from the stabilization of unfolded states, and played rather a role in stimulating folding.

GroEL thus displayed two interaction modes. In the first, unfolded substrates were bound, immobilized, and stabilized. In the second, they were compacted by attractive forces while preserving the necessary mobility to fold. Note that the compaction occurred at substantial forces, well above 10 pN (Fig. 2d, e). Such a collapse process, in which chains compact and form some secondary and tertiary structure, is considered key in spontaneous folding^25,34,35^. Two other substrates, dmMBP and rhodanese, displayed similar collapse enhancement by GroEL-ADP (Fig. 2e, Extended data Fig. 6). Notably, the slow folding dmMBP showed a large increase in *P_c_* from 0.06, to over 0.8 in presence of GroEL-ADP.

## Role of the GroEL apical domains

Next, we tried to disentangle the contributions of the GroEL apical domains to unfolded chain stabilization and folding. We performed MBP relax-wait-stretch experiments in the presence of GroEL-ADP, as it had yielded the highest *P*\*, but also added small peptides equivalent to the unstructured GroES loops, which compete strongly for the apical domain substrate binding sites^36^. Consistently, the sudden stabilization of unfolded chains (Extended data Fig. 2) was no longer observed, which is in line with reduced apical domain binding. Interestingly however, *P_c_* and *P*\* were now even higher than for GroEL-ADP only, both for MBP and dmMBP (Fig. 3, Extended data Fig. 7). These increases were not caused by the GroES-loops directly, since they alone did not yield increases (Fig. 3). These data are consistent with the apical domains antagonizing folding.

**Figure 3.**
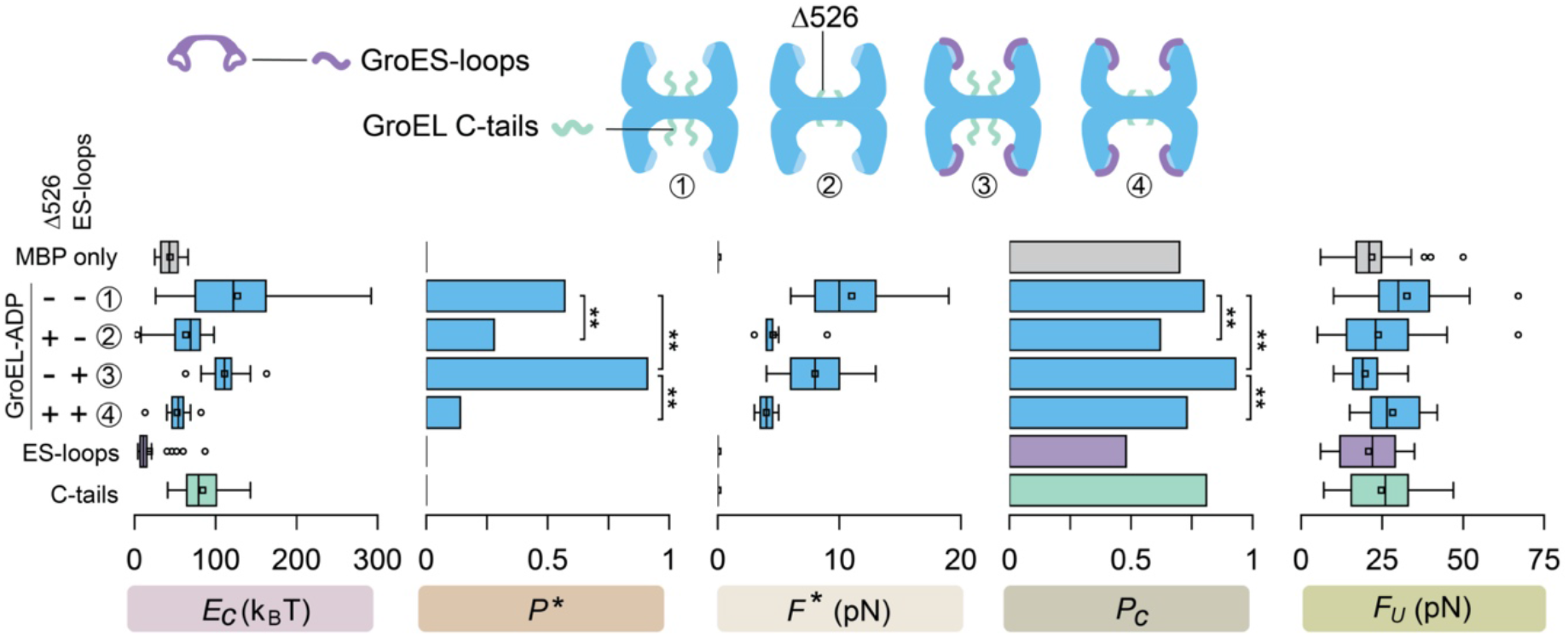
Roles of the GroEL apical domains and cavity. Top: Measured GroEL variants. The unstructured loops of GroES, added as separate polypeptides (purple), bind the apical domains (light blue) of GroEL and reduce substrate affinity. The effect of the unstructured C-terminal tails (light green) of GroEL is tested by truncation (GroELΔ526). Bottom: Parameter quantification from relax-wait-stretch cycles in different conditions (see Fig. 2b): total compaction energy during relaxation (*E_c_*), fraction of relaxation curves with steps (*P*\*) at force (*F*\*), core refold probability after 5 s. at 0 pN (*P_c_*), unfolding force or maximally sustained force (*F_U_*), as determined from MBP relax-stretch cycles. Conditions, from top to bottom: no chaperone (*N*=65), 200 nM GroEL and 1 mM ADP (*N*=96), 200 nM GroELΔ526 and 1 mM ADP (*N*=39) 200 nM GroEL, 1 mM ADP and 1 μM GroES loops (*N*=55), 200 nM GroELΔ526, 1 mM ADP and 1 μM loops (*N*=11), 1 μM GroES-loops only (*N*=29), and 1 μM C-tails only (*N*=32). Two stars: significant difference (*p*<0.05).

The apical domains also exhibited another effect: the observed unfolding force *F_u_* of MBP, which had in fact increased with GroEL-ADP, now decreased back to MBP-only levels when the GroES loops were present (22, 33, 23 pN respectively, Fig. 3, Extended data Fig. 8). dmMBP showed a similar trend (Extended data Fig. 7). These data indicated that GroEL-ADP can stabilize (partially) refolded states against forced unfolding, in addition to stabilizing unfolded states, while binding of the GroES-loops suppressed both effects. The finding that compaction and folding remained stimulated in the presence of GroES loops suggested that parts of GroEL other than the apical domain binding sites were relevant.

## Role of the GroEL C-terminal tails

To test whether the GroEL interior played a role in the folding stimulation, we truncated the unstructured C-terminal tails at the cavity bottom (GroELΔ526)^37^. *P_c_* and *P*\* were indeed more than two-fold lower for GroELΔ526-ADP than for GroEL-ADP, both with and without the GroES-loops present (Fig. 3, Extended data Fig.7). Interestingly, even alone the C-tails could promote some compaction (Fig. 3). Overall, the data showed that GroEL-mediated collapse and folding depended on the C-tails in the GroEL cavity. The enhancement of chain collapse while maintaining the dynamics that is required for folding upon interaction with GroEL is notable, and suggests a balance between different cavity properties^15,37^.

## The substrate-chaperone complex

Finally, we sought to verify two key interactions of the substrate-chaperone complex in these experiments, which required different approaches. To directly visualize GroEL-substrate binding, we scanned a fluorescence excitation beam along the tethered MBP during relax-wait-stretch cycles (Fig. 4a). ADP and Atto532-labeled GroEL were present at reduced concentrations to limit background fluorescence. The appearance of a fluorescent spot between the beads indicated binding of a single GroEL tetradecamer (Fig. 4b, yellow triangle). The tweezers measurements reported simultaneously on folding events (Fig. 4c, green triangle), in similar manner as shown previously (Fig. 2b). Consistently, during relaxation, such GroEL binding events always occurred first, and folding steps afterwards (Fig. 4d). These findings confirmed stimulated folding transitions in substrates complexed with GroEL.

**Figure 4.**
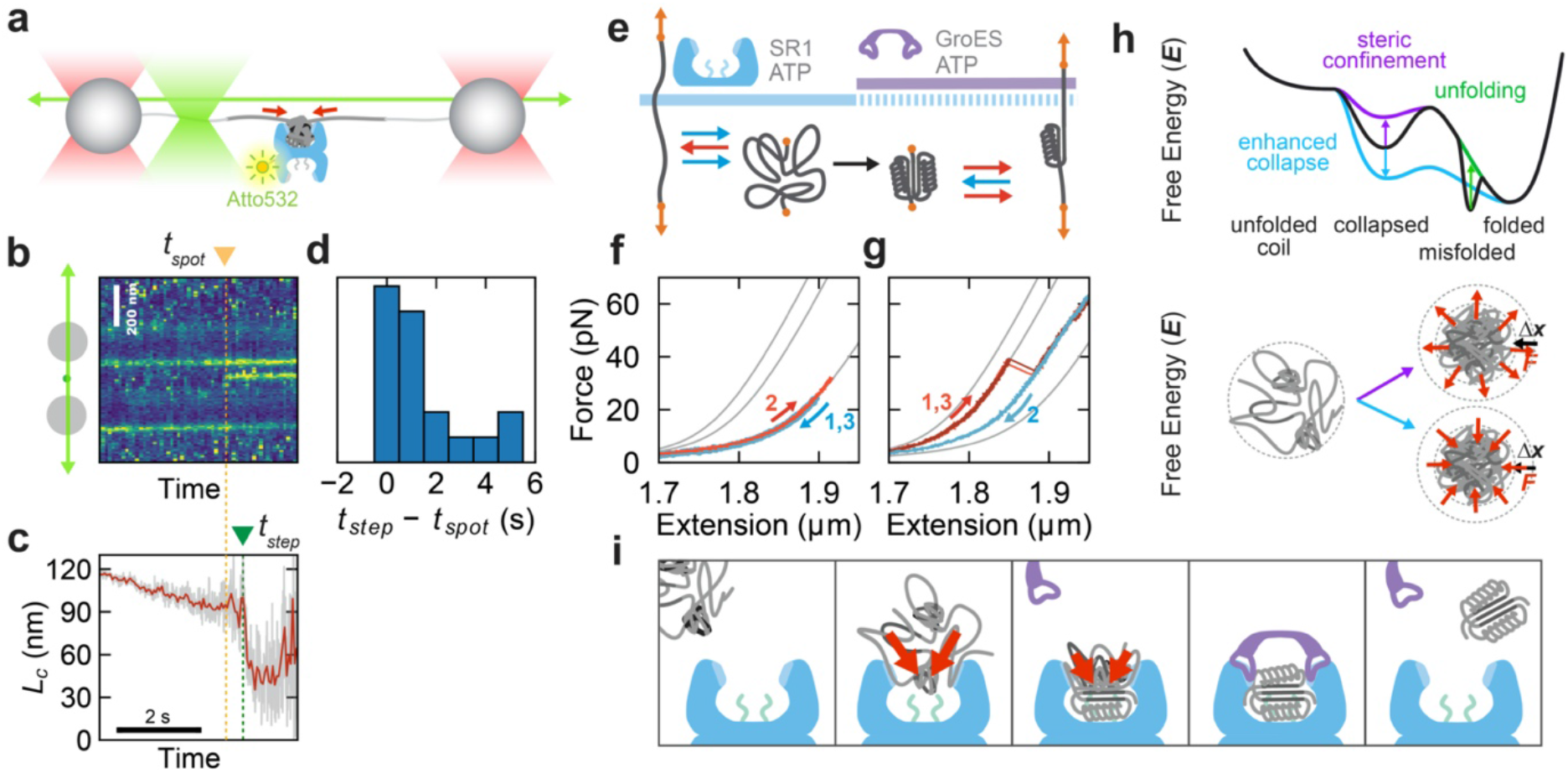
Compaction and folding in a single GroEL tetradecamer. **a**, Set-up to detect substrate-GroEL complex during stimulated folding, corresponding to panels b-d: optical tweezers (red beams) measure polypeptide compaction, while scanning fluorescence excitation (green beam) measures GroEL binding, in presence of GroEL (blue, 100 nM) labelled by Atto532 (yellow), and ADP (1 mM). **b**, Corresponding fluorescence emission kymograph during relaxation. Vertical lines show fluorescence from repeated scans along beads and substrate, during one relaxation experiment. Fluorescent spot at time *t_spot_* shows single GroEL binding. Beads also give a (constant) fluorescence signal, e.g. from GroEL adhesion. **c**, Measured contour length of non-compacted part of polypeptide (*L_c_*, red is filtered), shows step at time *t_step_*. **d**, Time between binding and step (*N* = 19). Times are positive, indicating GroEL binds first, and folding occurs afterwards when GroEL is bound. **e**, Buffer exchange protocol to test ternary complex (SR1-GroES-MBP) formation, corresponding to panels f and g. Buffer is exchanged by moving trapped beads to different chamber in flow cell. **f**, SR1 binding to substrate. In presence of 200 nM SR1 and 1 mM ATP, MBP remains unfolded during relaxation (1), stretching (2), and relaxation (3), consistent with binding to SR1 apical domains. **g**, GroES binding to SR1-substrate complex. After change to 500 nM GroES and 1 mM ATP, stretching data (1) follows MBP core WLC curve (middle gray trace), which unfolds partially during stretching. After relaxation (2), stretching shows core state again (3). Data are consistent with GroES displacing substrate from SR1 apical domains, followed by folding in SR1 chamber. **h**, Energy landscape cartoons for folding stimulation models. Green: GroEL exerts unfolding forces on misfolded states, raising their free energy to allow escape to other folded states. Purple: In steric confinement, chains are compressed (Δx) against counteracting entropic expansion forces, also raising their free energy, and effectively lowering the folding barrier. Blue: In enhanced collapse, chains are also compacted (Δx), but the mechanism differs: the contractile forces do not counteract but drive compaction, yielding a lower free energy. The folding barrier is lowered, as evidenced by the increased folding probabilities. **i**, Cartoons of event sequence suggested by our data. Substrate chain with folding-limiting autonomous collapse binds GroEL apical domains at different sites; GroEL-mediated forces draw unbound chain segments into the cavity bringing them closer together; stimulation of partial folding; competitive GroES binding releases bound segments, which are thus free to collapse and fold in stimulated manner; GroES and substrate are released. Various factors including substrate sequence, initial conformation, and chance encounters with apical domains may affect this process.

Second, we used a buffer-exchange protocol to verify that ternary complexes (GroEL-GroES-MBP) can indeed be formed in the optical tweezers assay. The idea was to employ the single-ring GroEL variant (SR1) for this purpose^38^. GroES binding to SR1 is known to trap the latter in the ADP-bound state with ATP present, and hence irreversibly lock GroES-SR1 together^13,14^. The binding of GroES to the SR1 apical domains is also considered to displace substrates from the apical domains into the SR1-GroES cavity.

The first step was to complex SR1 to unfolded MBP. Hence, we unfolded MBP in the presence of SR1 and ATP. Subsequent inability of MBP to refold during multiple relax-wait-stretch cycles indicated that SR1 had bound to the unfolded MBP chain (Fig. 4e, f). The beads that tethered MBP were then moved to another channel in the microfluidic device, which contained a medium with GroES and ATP only. Here, SR1 was not present to prevent premature binding of GroES to SR1. In this movement process, any unbound SR1 in solution was thus washed away, and only bound SR1 could be moved to the new measurement channel. This exposure to GroES and ATP was found to trigger MBP refolding to the core state, which in subsequent stretching did not unfold fully (Fig. 4g).

These data strongly suggested a SR1-ES-MBP complex: isolated and unbound core MBP states are known to unfold below 40 pN (Fig. 1c), while here it could not be unfolded fully even at 60 pN (Fig. 4g). Also note that GroEL was found to stabilize tertiary structure in similar manner (Fig. 2e), while MBP bound to SR1 (without GroES) was found to remain unfolded (Fig. 4f). In addition, the observed refolding upon exposure to GroES is consistent with the known role of GroES in competing substrates off the SR1 apical domains and displacing them into the SR1 cavity, where they subsequently fold. Because MBP has DNA linkers attached, these data suggested the SR1-GroES cavity is not hermetically sealed^39^. It is thus possible that GroES does not require intimate contact with all seven SR1 monomers to form a stable complex capable of initiating folding.

## Discussion

The folding stimulation by enhanced collapse that emerged from our experiments is distinct from current GroEL-ES models (Fig. 4h, i), while also presenting comparable features^13,14^. In confinement models, steric repulsion forces exerted by the walls of the GroEL cavity closed by GroES are proposed to decrease the chain entropy and thus increase its free energy, which effectively lowers folding barriers. In unfolding models, pulling forces on misfolds also increase the chain free energy and are proposed to allow escape to productive folding trajectories. Here, we measured the forces induced by GroEL-ES, and found that they are contractile, enhance protein chain collapse, and are mediated by open GroEL (unclosed by GroES). Such compaction suggests a decreased chain free energy, rather than an increased one (Fig. 4h). The observed increases in folding probabilities indicate that the folding barrier height was reduced (Fig.4h). Indeed, the spontaneous collapse of protein chains lowers the kinetic folding barrier by bringing residues together that must contact to form the folded structure^35^. Note that both confinement and collapse bring chain segments together, even as the compaction mechanism and resulting conformational dynamics differ. Hence, the model of enhanced collapse is consistent with diverse experimental findings^14,40^. Also in line with existing models, unfolded chain segments were observed to bind GroEL apical domains^14^, though those segments that did not bind, or were released by GroES were free to collapse, mediated by the contractile forces (Fig.4i). We note that collapse enhancement may well act in conjunction with steric confinement and misfold unfolding.

Our data thus provide a mechanism for the direct stimulation of protein folding, beyond promoting folding indirectly by limiting aggregation. While other direct stimulation mechanisms have been proposed (Fig. 4h), the required underlying compression and unfolding forces and resulting folding transitions have not been measured directly. The physical effects that underlie collapse enhancement may be similar to notions advanced in theoretical studies on the role of water within the GroEL-ES cavity^12^. More generally, charged and hydrophobic surfaces in particular in confined volumes, as well as amphiphilic solutes, can modulate the hydrophobic effect^41^, which in turn is the driving force in protein chain collapse. The collapse control by proteins observed here provides the possibility of local modulation and allosteric regulation of these effects.

GroEL-ES is known to support a broad range of proteins, and collapse is pervasive in unfolded proteins. One may speculate that non-optimal polypeptide collapse is a more general folding impediment that GroEL-ES helps resolve. Collapse enhancement by GroEL-ES can thus limit the lifetime of aggregation-prone collapsed states, and help drive the transfer of substrates from Hsp70 to GroEL. Moreover, we found that collapse modulation does not require the closed cavity that is unique to GroEL-ES. The ability to manipulate collapsed protein states may thus be exploited more generally within the protein quality control machinery to regulate folding, intrinsic disorder, and protein coacervation.

## Acknowledgements

Work in the group of S.J.T. is supported by the Netherlands Organization for Scientific Research (NWO). Work in the group of H.S.R was supported by a grant from the U.S. National Institutes of Health (R01GM114405). We thank A. L. Horwich, K. Chakraborty and B. Schuler for providing plasmids, and R. van Leeuwen, M. Mayer, J. van Zon, W. Noorduin, and P. R. ten Wolde for comments and critical reading of the manuscript

## Author Contributions

M.M.N and S.J.T. conceived and designed the research; M.J.A, E.J.K and V.S. designed and generated the substrate constructs; M.M.N. performed the optical tweezers experiments; A.R, H.S.R and G.K designed and purified the GroEL and GroES constructs; M.M.N, M.J.A, and S.J.T. analyzed the data; and M.M.N, M.J.A, H.S.R and S.J.T. wrote the manuscript.

## Methods

### Expression and purification of MBP, dmMBP and rhodanese

MBP and dmMBP were overexpressed in T7 competent cells (NEB laboratories) in LB medium supplemented with 0.2% glucose and 50 μg/ml kanamycin at 30°C until OD_600_~0.6, induced with 0.4 μM IPTG (Sigma) and incubated at 18°C overnight. The culture was harvested by centrifugation at 5000 g for 20 minutes at 4 °C. All following steps were carried out at 4 °C. The pellet was resuspended in ice-cold buffer A (50 mM phosphate buffer pH 7.5, 200 mM NaCl, 10 mM EDTA, 50 mM Glutamic Acid–Arginine (Sigma) and 3 mM ß-mercaptoethanol (Sigma)) and lysed using an Emulsiflex homogenizer. The lysate was cleared from cell debris by centrifugation at 50000 g for 1 hour followed by incubation with Amylose resin (NEB) for 1 hour. After extensive washing with buffer A, the proteins were eluted using buffer A supplemented with 20 mM maltose.

For rhodanese, the pellet was resuspended in buffer B (100 mM Tris-HCl pH 7.0, 5 mM EDTA, 20 mM Na_2_S_2_O_3_, 2 mM ß-mercaptoethanol) and lysed as described above. The lysate was mixed with Protino™ Ni-NTA Agarose (Macherey-Nagel) and incubated for 1 hour. After washing, the protein was eluted with buffer B supplemented with 250 mM imidazole.

### Purification of GroEL, GroES and their variants

GroEL was expressed from an inducible plasmid in *E. coli* BL21 in LB at 37 °C^1^. After cell disruption, the crude lysate was clarified by ultracentrifugation (142000 rcf), followed by anion exchange chromatography (FastFlow Q, GE) equilibrated in buffer C (50 mM Tris pH 7.4, 0.5 mM EDTA, 2 mM DTT) and eluted by linear gradient from 7.5% to 35% with buffer D (50 mM Tris pH 7.4, 0.5 mM EDTA, 2 M NaCl, 2 mM DTT). GroEL fractions were concentrated by 70% (w/v) ammonium sulfate precipitation. This precipitate was solubilized and dialyzed against 50 mM Bis-Tris pH 6.0, 50 mM KCl, 0.5 mM EDTA, 2 mM DTT containing 25% (wild-type GroEL) or 12.5% (all GroEL mutants) methanol. A second round of strong anion exchange (FastFlow Q, GE), run in the same methanol-containing buffer at pH 6.0, was used to strip co-purifying small proteins and peptides from the GroEL oligomers. To further remove contaminating proteins and peptides that remain tightly associated through prior stages of purification, GroEL fractions were gently agitated in the same methanol-containing buffer and Affi Blue Gel (BioRad) resin overnight at 4°C under an argon atmosphere. The final sample was dialyzed into storage buffer (25 mM Tris pH 7.4, 100 mM KCl, 0.5 mM EDTA, 2 mM DTT), supplemented with glycerol (15–20% v/v), concentrated, and snap frozen using liquid nitrogen.

GroES was expressed from an inducible plasmid in *E. coli* BL21(DE3) in LB at 37°C. After cell disruption, the crude lysate was clarified by ultracentrifugation (142,000 rcf), followed by acidification with sodium acetate, and cation exchange chromatography (FastFlow S, GE) equilibrated in buffer E (50 mM NaOAc pH 4.6, 0.5 mM EDTA, 2 mM DTT) and eluted by linear gradient from 0% to 25% buffer F (50 mM NaOAc pH 4.6, 0.5 mM EDTA, 2 M NaCl, 2 mM DTT). The sample was dialyzed against 25 mM Tris pH 7.4, 0.5 mM EDTA, 50 mM KCl, 2 mM DTT and applied to a strong anion exchange column (Source Q, GE). GroES was eluted with NaCl and enriched fractions were pooled. The sample was dialyzed into storage buffer supplemented with glycerol (15 – 20% v/v), concentrated, and snap frozen using liquid nitrogen.

For the expression of Single Ring GroEL (SR1), *E. coli* BL21 DE3 transformed with pSR1 was grown in LB-Ampicillin (100 μg/ml) at 30 °C to an OD_600_=0.5. Overexpression was induced by adding 1 mM IPTG and growth was continued for 3 hours. Cells were harvested by centrifugation and stored at −70 °C after flash freezing in liquid nitrogen. Frozen cells were resuspended in 20 mM Tris-HCl pH 7.4, 50 mM KCl, 1 mM EDTA, 1 mM DTT, 1 mM PMSF, lysed using a French Press and cell debris were removed by centrifugation. SR1 was enriched by fractionated (NH_4_)_2_SO_4_ precipitation between 35% and 45% saturation. Following dialysis in 50 mM Tris-HCl pH 8, 1 mM EDTA at 4 °C, the protein solution was fractionated using a DEAE Sepharose Fast Flow anion exchange chromatography resin (GE healthcare) eluting with a gradient from 0 to 1 M NaCl and further fractionated by size-exclusion chromatography using a HiPrep™ 26/60 Sephacryl^®^ S-500 HR column. SR1 containing fractions were pooled, concentrated using Amicon Ultra centrifugal filters (Merck), frozen in liquid nitrogen and stored at −70 °C.

### GroEL labeling

The GroEL variant (EL315C)^2^ was labeled with Atto-532 maleimide (Sigma). Reactive dyes were prepared fresh from dry powder in anhydrous dimethylformamide (DMF) immediately prior to use. All proteins were first buffer exchanged 300-400x the original volume by a Vivaspin Turbo 15 (Satorious) into 50 mM Tris buffer pH 7.4, 100 mM KCl, 0.5 mM EDTA, and 1 mM TCEP. The proteins were then run over gel filtration (PD-10 column; Pharmacia) equilibrated in reaction buffer (50 mM Tris pH 7.4, 100 mM KCl, 0.5 mM EDTA, 0.5 mM TCEP). EL315C was concentrated to a final concentration of 70 μM (monomer) in a volume of 5 mL. Protein samples were added to individual 5 mL conical Weaton reaction vials, followed by two sequential reactive dye additions. Freshly prepared Atto-532 maleimide in DMF was added at a molar ratio of 1:6.5 to EL315C monomer. Following each addition, the sample was incubated for 45 minutes in the dark at 23°C. Following the full 1.5 hours reaction time, the sample was quenched by addition of 5 mM glutathione. The labeled EL315-Atto532 were separated from unreacted dye by four rounds of dilution and concentration in a Vivaspin Turbo 15 (Sartorious), followed by gel filtration (PD-10 column; Pharmacia). The labeled proteins were then supplemented with glycerol (15-20%) and snap frozen using liquid nitrogen. Protein concentration was determined using a calibrated Bradford assay, in which the protein standard was from a sample of wild-type GroEL whose concentration had been previously established. Conjugated dye concentrations are determined by absorption spectroscopy of the denatured proteins (in 6 M Gdm buffer) using the following corrected extinction coefficient: Atto-532, 115000 M^−1^cm^−1^. GroEL-Atto 532 activity was confirmed by MESG ATPase activity assay (EnzChek, Molecular Probes) and native gel filtration (Superdex 200, GE).

### GroES mobile loops and GroEL C-tails

The GroES mobile loops^3^ ETKSAGGIVLTGS and GroEL C-tails (GGM)_4_M were ordered from Genscript. GroES mobile loops and C-tails were dissolved in MQ water and snap frozen using liquid nitrogen. Prior to measurements the samples were dissolved in HMK buffer (50 mM HEPES, pH 7.5, 5 mM MgCl2, 100 mM KCl). The GroES mobile loops were added in fivefold molar excess to GroEL during optical tweezers experiments (Fig. 3, Extended data Fig. 7).

### Protein-DNA constructs

The cysteines at the N and C termini of proteins were coupled with 20 bp maleimide ssDNA oligos at 37°C for one hour. 2.5 kbp and 1.3 kbp DNA tethers were generated by PCR from pUC19 plasmid (NEB) with a double digoxigenin- or biotin-labeled primer on one side and a phosphoprimer on the other side. Purification was done with the QIAquick PCR purification kit (Qiagen). The phosphorylated strand was digested by Lambda exonuclease (NEB) for 2 hours at 37°C and purified using a Amicon 30 kDa MWCO filter (Merck). Deep Vent exo-DNA polymerase (NEB) and a 20 nt more upstream primer than the phosphoprimer from the PCR was used for the fill up of the second DNA strand creating a 20 nt overhang. This overhang is complementary to the 20 nt oligonucleotide sequence coupled to the termini of proteins. The overhang DNA was added to the protein-oligo chimera together with T4 ligase (NEB) and incubated for 30 min at 16°C followed by 30 min on ice. The resulting protein-DNA hybrid was flash frozen and stored at −80°C until measurement.

### Optical tweezers experiments

Neutravidin coated beads (2.1 μm) were purchased from Spherotech and stored at 4°C until use. Anti-digoxigenin beads were prepared by coating carboxylated polystyrene beads (2.1 μm, Spherotec) with anti-digoxigenin antibodies from Sigma-Aldrich using a carbodiimide reaction (Poly-Link Protein Coupling Kit, Polysciences Inc.) The protein coated beads were prepared by mixing 50 ng of MBP, dmMBP or rhodanese constructs with anti-digoxigenin beads in 10 μl HMK buffer. The mixture was then incubated at 4 °C for 30 min on a rotary mixer. Next, the beads were dissolved in 400 μl HMK buffer for optical tweezers experiments. Optical tweezers measurements were done in HMK buffer. ADP and ATP solutions were prepared by dissolving ADP and ATP sodium salt from Sigma Aldrich in HMK buffer. Experiments in GroEL ADP conditions were verified using ultra-pure ADP (99.9%, Gentaur).

Stretch-relax experiments were performed on two optical tweezers setups. The first was a custom-built single trap instrument. A substrate-coated anti-digoxigenin bead was held in the optical trap and a NeutrAvidin bead was placed on the end of a micropipette tip. The two beads were brought in close contact, allowing a tether between the beads to form. Proteins were stretched and relaxed by moving the flow-cell and micropipette with a nanopositioning piezo stage at 50 nm/s speed which corresponds to a pulling rate of 5 pN/s. The deflection of the bead in the trap was measured using quadrant photodiode at 50 Hz. The data were filtered with a 5th order Butterworth filter at 20 Hz. The optical traps were calibrated by recording the power spectrum of the Brownian motion of the beads^4^ yielding stiffnesses ranging from 120 – 170 pN/μm.

The second setup was a dual trap optical tweezers instrument (C-trap from Lumicks). As described above, tethers were formed by bringing similarly prepared construct-coated and NeutrAvidin beads in close proximity. The protein was stretched and relaxed at a constant velocity of 50 nm/s, by moving one of the traps. The data was acquired at 500 kHz and averaged to 500 Hz. For constant force measurements, tension was held at 2 pN on average for 30 s using a proportional–integral–derivative (PID) feedback loop, before pulling again at constant velocity (Fig. 1e). In Extended data Fig. 2c, the distance between the traps is constant, while the extension of the protein is monitored as it changes conformation. Note that the beads can change position within the traps. For fluorescence measurements in combination with stretch-relax experiments (Fig. 4a to d), Atto-532 labelled GroEL proteins were visualized using a green excitation laser (532 nm), with 2 mM Trolox and 4 mM ß-mercaptoethanol in the buffer. The excitation beam was used to scan along the tethered construct at 10 Hz during the force-spectroscopy measurements, generating fluorescence kymographs that were aligned to the force signal using ImageJ.

### Data Analysis

Several checks were performed to confirm that the data corresponded to a proper single tether, which include comparing the total measured unfolded length to the expected length, consistency with the WLC model (at higher forces), overstretching at 67 pN, and final tether breakage in one clean step. The unfolding forces (*F_U_*), contour lengths (*L_C_*), refolding forces (*F*\*) and compaction energies (*E_C_*) were quantified from force extension data, using an open source MATLAB code^5^ after modifications. *F_U_* was determined from stretching traces as the force required to fully unfold a protein (Fig. 1c left, e, 2c to e, 3, Extended data Fig.7). For stretching traces in which the protein did fully unfold below the maximum force that could be applied (67 pN, corresponding to the DNA overstretching plateau), *F_U_* was determined as 67 pN, the maximally sustained force (Extended data Fig. 8, Fig. 2e). The contour lengths (*L_c_*) of refolded states were determined from the force-extension data of the stretching curve before the first unfolding transition, using the WLC model (Extended data Fig. 1a, b). The persistence lengths of the DNA (45 nm) and protein (1.5 nm), and the stretch modulus of DNA (1200 pN) were fixed parameters in the WLC model. In Fig 2a and 4c, the instantaneous protein contour length was calculated using the same WLC model. Compaction energy (*E_C_*) was calculated by quantifying the area under the relaxation curve and then subtracting the area under the WLC curve for fully unfolded protein (Extended data Fig. 4b, c, 7, Fig. 2e). *P*\* was determined as the fraction of relaxation traces that show (one or more) steps in *L_c_* of more than 15 nm (Fig. 2e, 3, Extended data Fig. 7). *F*\* was quantified as the measured force just before such a step in *L_c_* (during relaxation, Fig. 2e, 3, Extended data Fig. 7). The folding probability (*P_c_*) was quantified as the fraction of relax-stretch cycles showing refolding to the core MBP state (Fig. 2e, 3, Extended data Fig. 1, 7).

### Statistical Analysis

The statistical significance of differences in folding probability (*P_c_*) and refolding at force probability (*P*\*) between experimental conditions was calculated using one tailed two proportion z-test. The statistical significance of differences in compaction energy (*E_c_*) and maximally sustained forces (*F_U_*) between experimental conditions was calculated using two sample assuming unequal variance t-Test. Test results are mentioned as *p* values in the main text. In box charts, whiskers indicate 90% and 10% extreme values, the inner line represents the median, the length of the box indicate interquartile range and the inner small square the mean of the population.

### Data availability

All data supporting the findings of this study are available in the main text and extended data figures. The raw data that support the findings of this study are available from the corresponding authors upon reasonable request.

### Code availability

The MATLAB code used for analysis is available from the corresponding author upon reasonable request.

## Extended Data Figures

**Extended data Figure 1.**
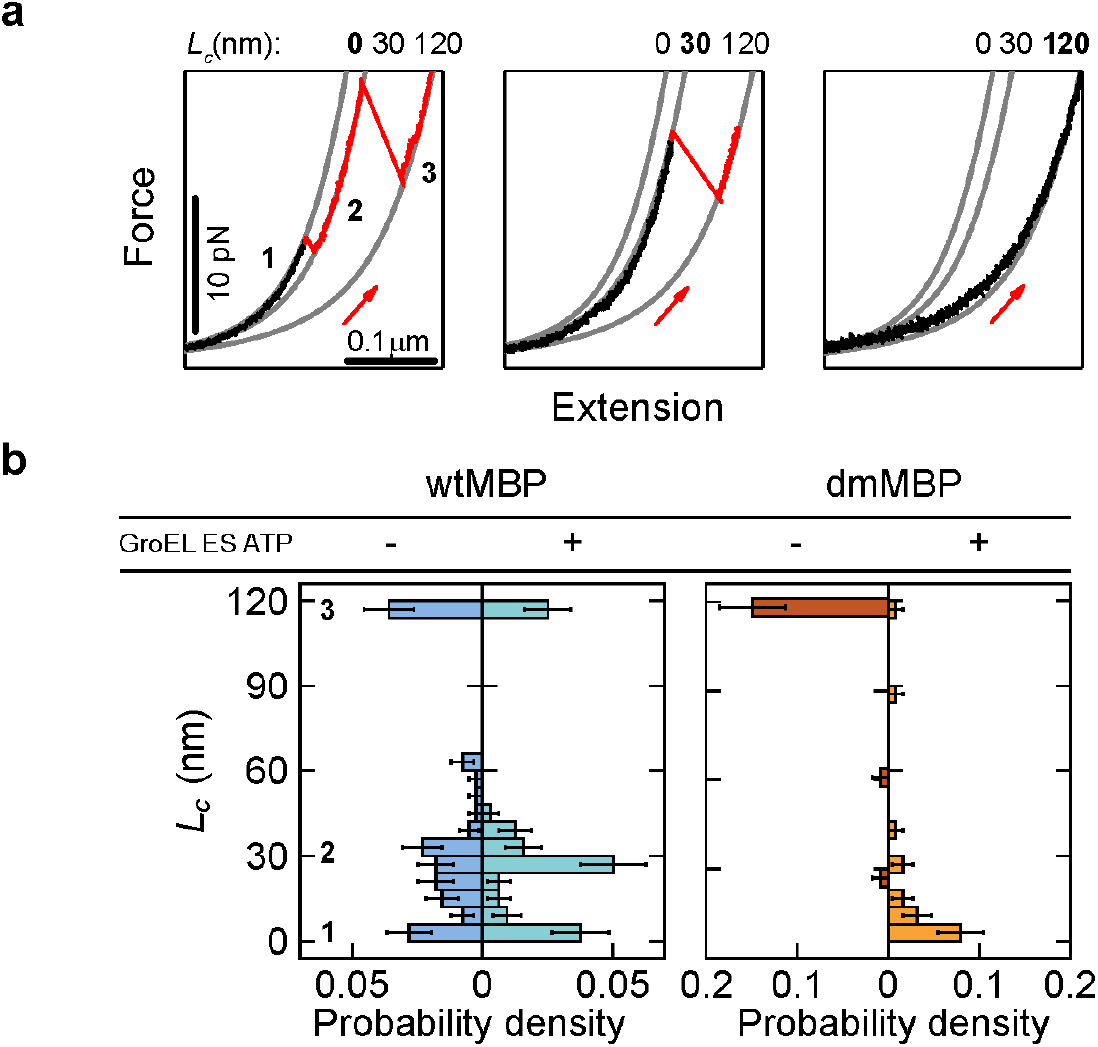
Contour lengths of refolded states. **a**, Determination of contour lengths (*L_c_*) from force-extension data taken during stretching. Displayed stretching curves are taken after relaxation of unfolded chains and waiting for 5 s at 0 pN to allow refolding. The protein states after this refolding window are characterized by their contour lengths (*L_c_*), which represents the length of the non-compacted part of the protein chain. This *L_c_* of refolded states is determined based on the first part of the stretching data, where no (detectable) unfolding has yet occurred (in black). This data is described by a single mean *L_c_* value that is determined using the worm-like chain (WLC) model (gray curves). Gray curves: force-extension behavior of DNA tethers attached to an unfolded protein chain of three different contour lengths *L_c_*. Indicated are WLC curves for fully compacted (*L_c_*=0 nm), MBP core state (*L_c_* = 30 nm), and fully unfolded state (*L_c_* = 120 nm). Panels indicate different example stretching curves that are observed for MBP with 200 nM GroEL, 500 nM GroES, and 1 mM ATP. Left panel: refolded state *L_c_* = 0 nm (1), followed by unfolding to core state (2), and to fully unfolded state (3). Middle panel: refolded state *L_c_* = 30 nm (core state), which then unfolds to fully unfolded state. Right panel: refolded state *L_c_* = 120 nm (fully unfolded state). For traces that show no unfolding transitions, like the latter example, *L_c_* is measured at 10 pN. **b**, Probability density (P.D.) of contour lengths (*L_c_*) of refolded states. Determination of *L_c_* as described in panel a, for MBP (blue), and dmMBP (orange) in the presence and absence of GroEL-ES and ATP. This analysis is also performed to quantify the frequencies (*P_c_*) of refolding to the core MBP structure. Also note that refolded states larger than the core structure always displays unfolding to (via) the core state, which involves detachment of MBP c-terminal helices from the core structure. Error bars are s.d.

**Extended data Figure 2.**
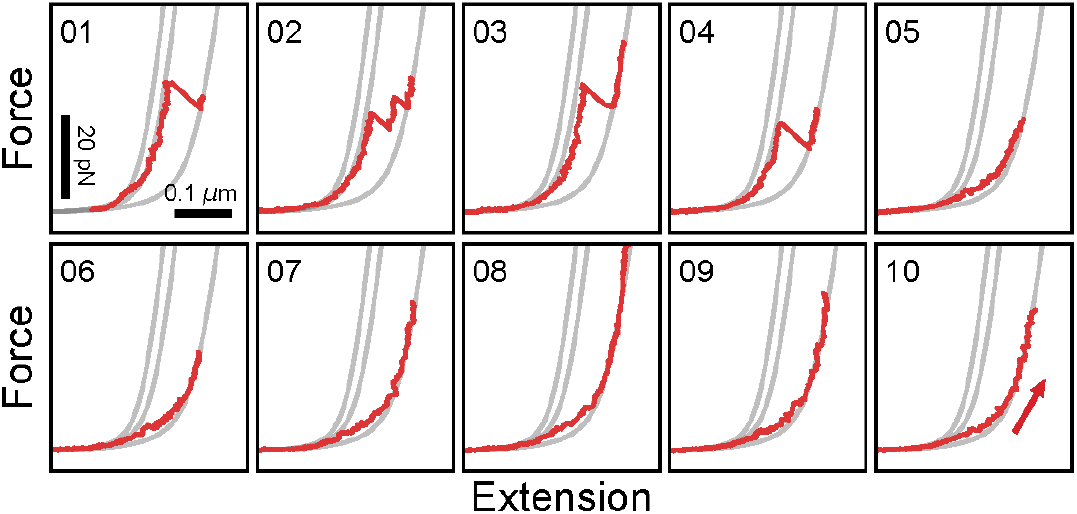
Irreversible switching to unfolded states in presence of GroEL. Force extension traces of MBP showing a sudden switch to a stable unfolded state. Successive stretching traces from relax-stretch cycles for MBP in the presence of 200 nM GroEL and 1 mM ADP, which initially (cycles 1 to 4) show the data following the worm-like chain (WLC) curve of the MBP core state (middle gray line) and thus indicating core refolding, followed by progressive unfolding to the fully unfolded state (right gray line). However, the data follows the WLC curve of the unfolded state and shows no unfolding transitions in the subsequent cycles (5 to 10), indicating the chain remained stabilized in the unfolded state. The data was close to the WLC curve corresponding to the unfolded state, though the deviation at lower forces suggested that a compacted yet non-folded state continued to be formed and disrupted.

**Extended data Figure 3.**
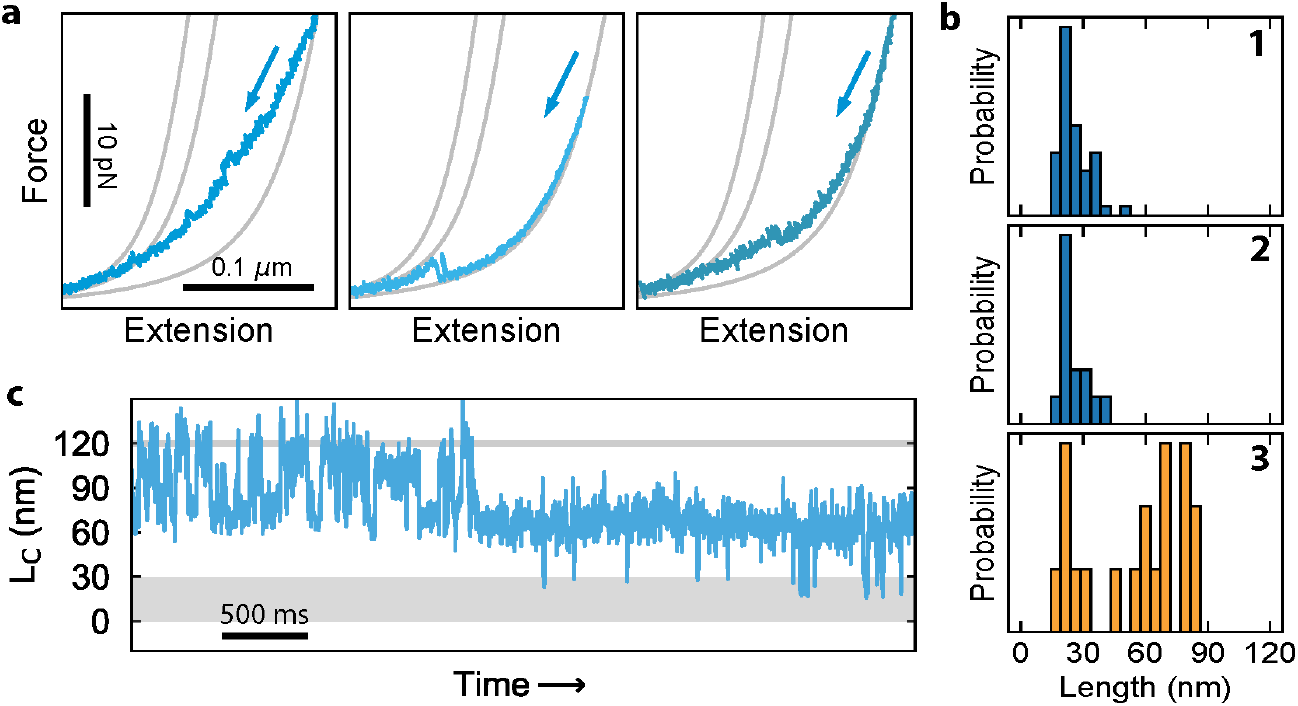
GroEL-mediated chain compaction and hopping transitions. **a**, Three example relaxation curves, displayed in force-distance graph, showing gradual and step-wise compaction, for dmMBP in the presence of 200 nM GroEL and 1 mM ADP condition. **b**, Histograms of contour length changes of step-wise compactions during relaxation (see panel a) for MBP, 200 nM GroEL and 1 mM ADP (1, N = 52), MBP, 200 nM GroEL398 and 1mM ATP (2, N = 14) and dmMBP, 200 nM GroEL and 1 mM ADP (3, N = 19) conditions **c**, Contour length vs time trace, showing repeated step-wise transitions (hopping) between states for dmMBP, mediated by GroEL. Data is taken with both traps at a constant position, in the presence of 200 nM GroEL and 1mM ADP. We stress that the details of these data, such as the folding step-sizes, are specific to this experiment and not generally observed. The latter may be expected. For hopping transitions of isolated proteins without chaperones, the energy landscape is defined only by the tethered protein that is in principle the same for different experiments. In contrast, here the energy landscape is also defined by GroEL, and how and where it is interacting with the substrate, which has a random aspect, and hence will produce differences between experiments.

**Extended data Figure 4.**
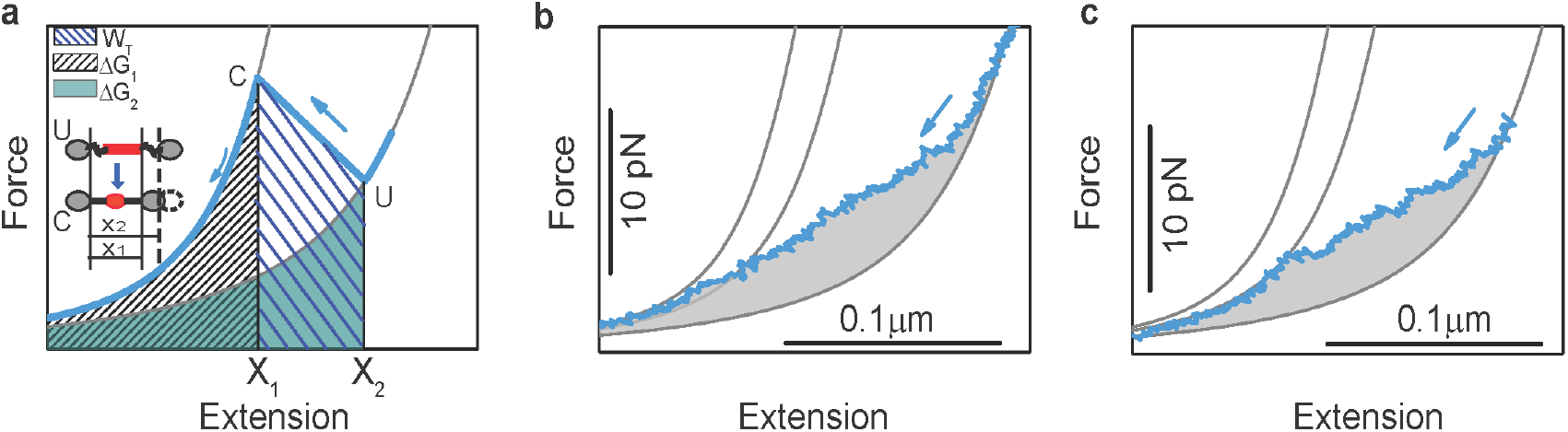
Determination of compaction energy. **a**, Schematic diagram showing how the compaction energy (*E_c_*) is determined from force extension cycles. We consider an unfolded protein chain during relaxation that suddenly compacts fully, but the derivation can also be used for gradual compaction composed of multiple smaller compaction events. At this single sudden event (U->C), the measured distance between beads (Extension, along the x-axis) suddenly *decreases* from x_2_ to x_1_, and the measured force *F*(*x*) that acts on these beads and throughout the DNA-protein-DNA tether (Force, along the Y-axis) *increases* from *F*(*x_2_*) to *F*(*x_1_*), because the tether is now effectively shorter and hence its tension higher (see inset in panel a). In the case of slow relaxation where the system is in equilibrium and there is no heat dissipation, energy is conserved, and hence the corresponding increase in potential energy of the beads equals to the work done (*W_T_*) by DNA-protein-DNA tether. *W_T_* is then estimated as the area under the force-extension curve from *x_2_* to *x_1_* that quantifies the increase in bead potential energy. Note that displacing an object over distance *dx* against a force *F* costs an amount of energy *F·dx*. Thus, *W_T_* is quantified by 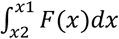 (wide-hash region). *W_T_* can be decomposed into two contributions: the work done to compact the protein (*E_c_*) and to increase the tension in the DNA-protein-DNA tether (*W_ext_*), hence: *W_T_* = *W_ext_* + *E_c_*. *W_ext_* can be calculated using the WLC model. To use this, we write *W_ext_* = ΔG_2_ – ΔG_1_, where ΔG_2_ is the work done in extending the unfolded protein chain and DNA linkers from extension 0 to x_2_ (green region), which is calculated with the WLC model. ΔG_1_ is the work done in extending only DNA from 0 to x_1_ (narrow-hash region), and is also calculated with the WLC model. ΔG_1_ has no contribution from the protein chain because it is fully compacted in this state. The work done in compacting the protein (*E_c_*) can then be calculated as *W_T_ + ΔG_1_ – ΔG_2_*. Thus, in graphical terms, *E_c_* equals the size of the wide-hash (*W_T_*) plus narrow-hash (*ΔG_1_*) regions minus the size of the green region (ΔG_2_). In more simple terms, this is thus the size of the area under the measured curve *F*(*x*) minus the size of the area under the WLC curve for the unfolded protein (right gray curve), as illustrated for measured data in panels b and c. Note that in the latter one can integrate from *x* = 0 to any *x* > *x_2_*, as beyond *x_2_* the chain is fully unfolded and hence there are no further area contributions. Perhaps counterintuitively, *E_c_* is thus determined not only by *F*(*x*) for *x* in between x_1_ and *x_2_*, but also by *F*(*x*) for *x* in between 0 and *x_1_*. Note that while the compacted chain (C) may in principle be deformed for *x* < *x_1_*, the length changes as well as energies are negligible, owing to its high stiffness compared to the DNA, while the force is identical throughout the chain. Finally, we note that this estimate of *E_c_* is a lower-bound, given that the system is not fully in equilibrium. (**b to c**) For gradual collapse and collapse with refolding jumps, the compaction energy *E_c_* is thus determined by the size of the indicated gray area.

**Extended data Figure 5.**
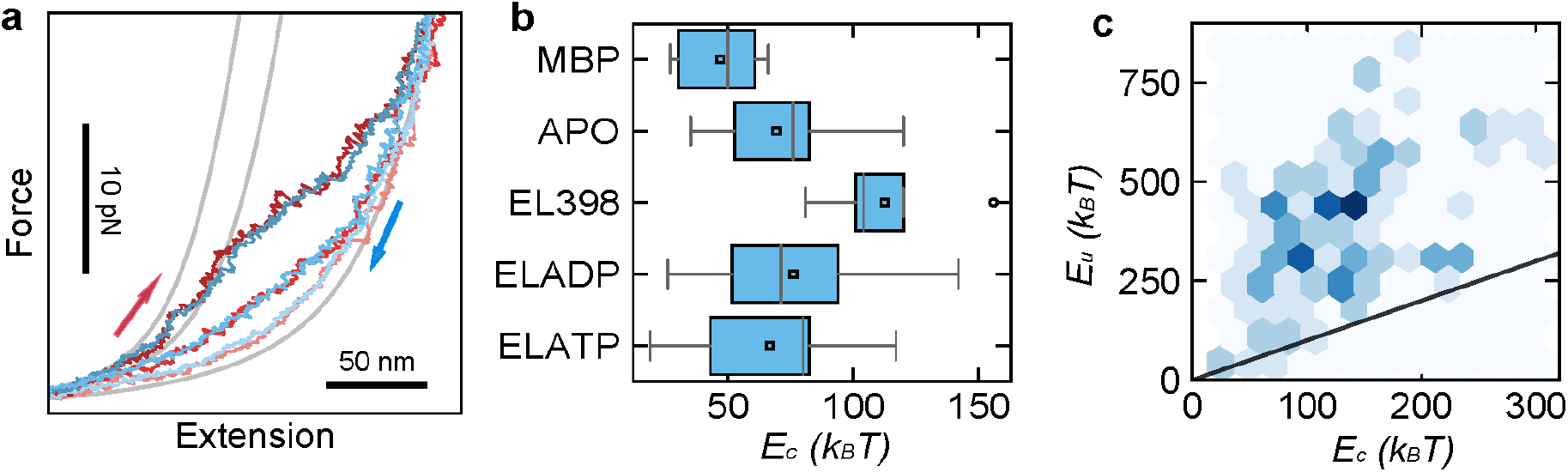
Collapse without folding, and stretching energy. **a**, Force-Extension traces that show compaction during relaxation (blue), and after waiting at 0 pN for 5 s, are followed by stretching (red) that do not show unfolding features. These cycles indicate that significant compaction can occur without the formation of detectable folded states. Note that most cycles in these conditions rather show refolding instead (See Fig. 2 main text). The displayed data is for MBP in presence of 200 nM GroEL and 1 mM ADP. **b**, Corresponding compaction energy (*E_c_*) of relaxation traces that do not produce detectable refolding (for examples see panel a) in different conditions. *E_c_* is highest for EL398ATP and ELADP (as in Fig. 2e). **c**, Area under the stretching curve (stretching energy *E_u_*) against the compaction energy (*E_c_*) from the prior relaxation trace for MBP with 200 nM GroEL and 1mM ADP condition. The data here includes both cycles that do show refolding and cycles that do not. Black line indicates *E_u_* = *E_c_*. Relax-stretch cycles producing no folding (panels a and b) are close to this line.

**Extended data Figure 6.**
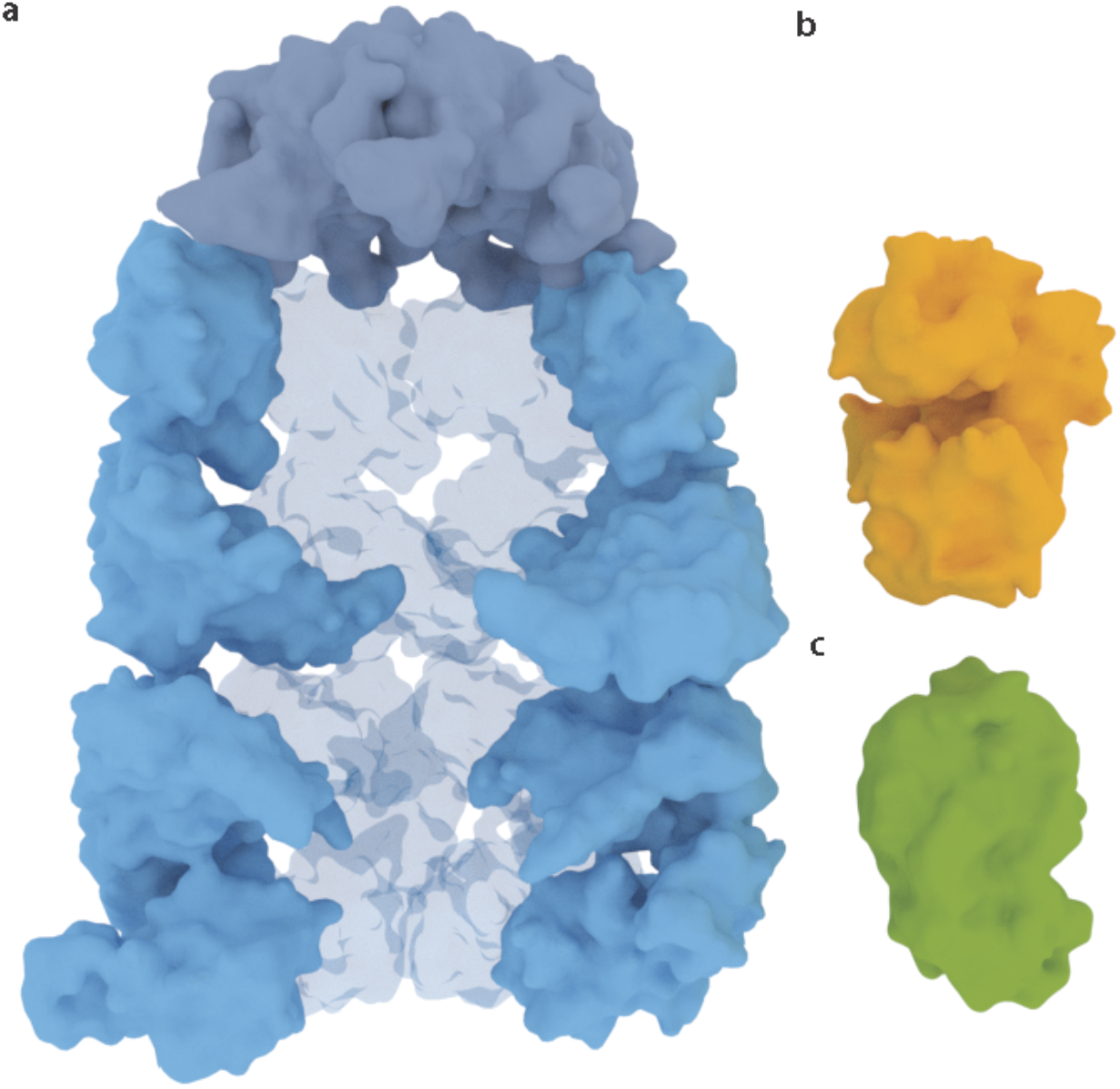
Protein structures. **a**, A medial slice of GroEL-GroES ADP bullet, side view (PDB: 1PF9). **b**, MBP (PDB ID: 2MV0) in orange. **c**, Rhodanese (PDB ID: 1rhs) in green. Proteins are displayed in the same scale to compare their relative sizes.

**Extended data Figure 7.**
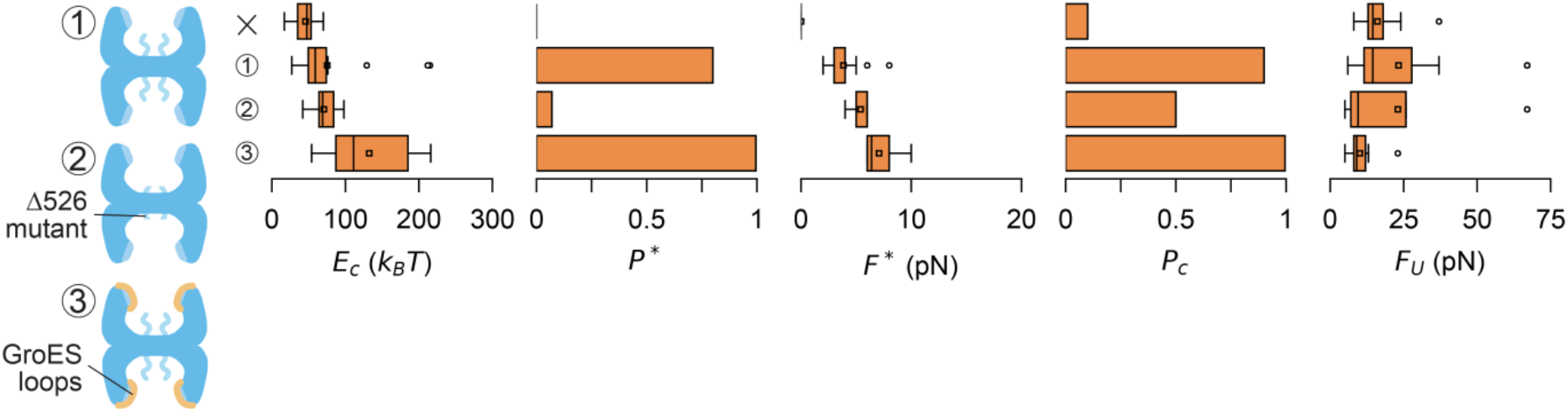
Roles of GroEL apical domains and cavity, dmMBP data. From dmMBP relax-stretch cycles, we quantified: total compaction energy during relaxation (*E_c_*), probability (*P*\*) at force (*F*\*) of steps during relaxation, core refold probability after 5 s. at 0 pN (*P_c_*), unfolding force (*F_U_*). Conditions are, from top to bottom: No Chaperone (x, *N*=19), 200nM GroEL and 1mM ADP (1, *N*=21), 200nM ELΔ526 and 1mMADP (2, *N*=11) and 200nM GroEL, 1mM ADP and 1μM loops (3, *N*=16). For dmMBP without GroEL (x), *F_U_* of first pulls is displayed because of the low refolding rate.

**Extended data Figure 8.**
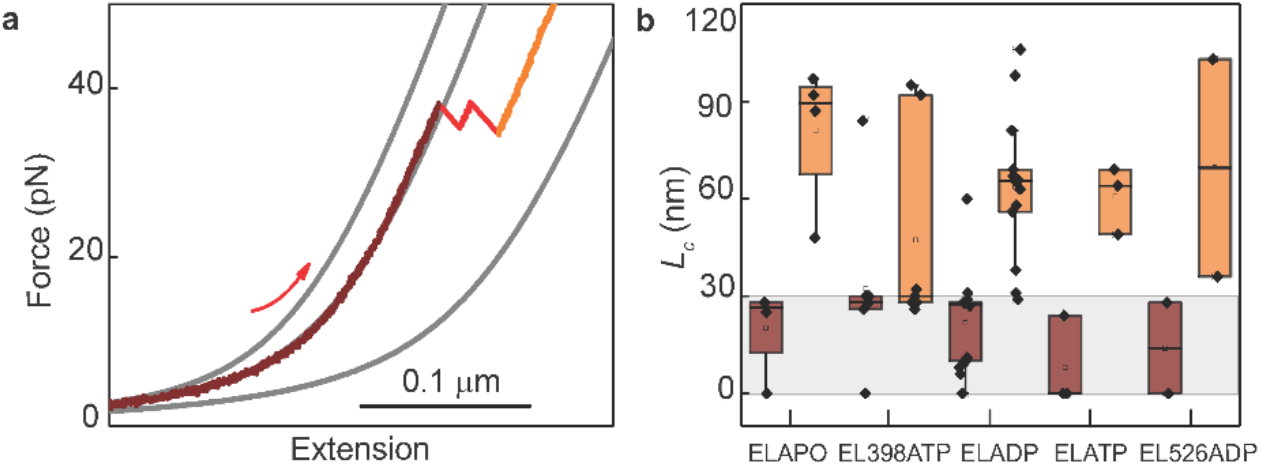
Lengths of stabilized partially folded structures. **a**, Stretching curve showing stabilization of partially folded MBP states against forced unfolding, in the presence of GroEL-ADP. Displayed data initially follows worm-like chain (WLC) curve of MBP core state (dark red), then unfolds partially in two steps (red), to a partially folded state that is stable against high applied forces (orange). **b**, Distributions of contour lengths *L_c_*. Dark orange: *L_c_* of initial MBP stretching data (panel a, dark red). Light orange: *L_c_* of observed MBP structures that are stable against forces over 40 pN (panel a, orange). 40 pN is the maximum force in the absence of chaperonin; see Fig. 2 e main text. The data indicates that the GroEL-stabilized MBP structures are typically smaller than the core state, in different nucleotide conditions.

**Extended data Table 1.** Experimental conditions. For each figure, the used substrate, chaperone components, nucleotides, and number of relax-wait-stretch cycles are indicated.

| Figure | Substrate | Chaperone components & nucleotides | N |
| --- | --- | --- | --- |
| 1c | MBP | no chaperones | n.a. |
| 1d | MBP | no chaperones | 65 |
|  | MBP | 200 nM GroEL, 500 nM GroES, 1 mM ATP | 53 |
|  | MBP | no chaperones | 18 |
|  | MBP | 200 nM GroEL, 500 nM GroES, 1 mM ATP | 21 |
|  | dmMBP | no chaperones | 19 |
|  | dmMBP | 200 nM GroEL, 500 nM GroES, 1 mM ATP | 21 |
| 1e | MBP | no chaperones | 65 |
|  | MBP | 200 nM GroEL, 500 nM GroES, 1 mM ATP | 53 |
|  | MBP | no chaperones | 18 |
|  | MBP | 200 nM GroEL, 500 nM GroES, 1 mM ATP | 21 |
|  | dmMBP | no chaperones | 19 |
|  | dmMBP | 200 nM GroEL, 500 nM GroES, 1 mM ATP | 21 |
| 2a | dmMBP | no chaperones | n.a. |
|  | dmMBP | 200 nM GroEL, 1 mM ADP | n.a. |
| 2e | MBP | no chaperones | 65 |
|  | MBP | 200 nM GroEL | 33 |
|  | MBP | 200 nM GroEL398, 1 mM ATP | 43 |
|  | MBP | 200 nM GroEL, 1 mM ADP | 96 |
|  | MBP | 200 nM GroEL, 1 mM ATP | 21 |
|  | dmMBP | no chaperones | 19 |
|  | dmMBP | 200 nM GroEL, 1 mM ADP | 21 |
|  | rhodanese | no chaperones | 16 |
|  | rhodanese | 200 nM GroEL, 1 mM ADP | 20 |
| 2b | MBP | 200 nM GroEL, 1 mM ADP | n.a. |
| 2c | MBP | 200 nM EL398, 1 mM ATP | n.a. |
| 2f | MBP | 200 nM GroEL398, 1 mM ATP | 43 |
|  | MBP | 200 nM GroEL, 1 mM ADP | 21 |
| 3 | MBP | no chaperones | 65 |
|  | MBP | 200 nM GroEL, 1 mM ADP | 96 |
| | MBP | 200 nM GroELΔ526, 1 mM ADP | 39 |
| | MBP | 200 nM GroEL, 1 μM GroES loops, 1 mM ADP | 55 |
| | MBP | 200 nM GroELΔ526, 1 μM GroES loops, 1 mM ADP | 11 |
| | MBP | 1 μM GroES loops | 29 |
| | MBP | 1 μM GroEL C-terminal tails | 32 |
| 4b | MBP | 100 nM GroEL-atto532, 1 mM ADP | n.a. |
| 4c | MBP | 100 nM GroEL-atto532, 1 mM ADP | n.a. |
| 4d | MBP | 100 nM GroEL-atto532, 1 mM ADP | 19 |
| 4f | MBP | 200 nM SR-1, 1 mM ATP | n.a. |
| 4g | MBP | 500 nM GroES, 1 mM ATP | n.a. |

